# CD44 facilitates adhesive interactions in airineme-mediated intercellular signaling

**DOI:** 10.1101/2024.02.27.582398

**Authors:** Raquel L Bowman, Jiyea Kim, Michael J Parsons, Dae Seok Eom

**Affiliations:** Department of Developmental and Cell Biology, University of California, Irvine, CA 92617, USA; Center for Complex Biological Systems, University of California, Irvine, CA 92697, USA; UC Irvine Skin Biology Resource Center, University of California, Irvine, CA 92697, USA

## Abstract

Specialized cellular protrusions facilitate local intercellular communications in various species, including mammals. Among these, airinemes play a crucial role in pigment pattern formation in zebrafish by mediating long-distance Notch signaling between pigment cells. Remarkably, airinemes exhibit large vesicle-like structure at their tips, which are pulled by macrophages and delivered to target cells. The interaction between macrophages and Delta-ligand carrying airineme vesicles is essential for initiating airineme-mediated signaling, yet the molecular detail of this interaction remains elusive. Through high-resolution live imaging, genetic *in vivo* manipulations and *in vitro* adhesion assay, we found that adhesive interactions via the extracellular domain of CD44, a class I transmembrane glycoprotein, between macrophages and airineme vesicles are critical for airineme signaling. Mutants lacking the extracellular domain of CD44 lose their adhesiveness, resulting in a significant reduction in airineme extension and pigment pattern defects. Our findings provide valuable insights into the role of adhesive interactions between signal-sending cells and macrophages in a long-range intercellular signaling.

## Introduction

Proper signal delivery between cells is essential in development and homeostasis in multicellular organisms. Even single-celled organisms communicate with each other for their survival in specific environments (Combarnous and Nguyen, 2020). Consequently, understanding the mechanisms of intercellular signaling has been a central topic in biology and medicine for decades. Although several cell-to-cell communication modalities have been identified, recent advancements in microscopy and techniques enabled us to uncover previously unappreciated mechanisms across various species and contexts. Specialized cellular protrusions or also known as signaling filopodia are one of them. These are long, thin cellular extensions similar to neuronal axons or dendrites but are present in non-neuronal cells, spanning from sea urchins to mice *in vivo* (Daly et al., 2022; Hall et al., 2024; Kornberg and Roy, 2014a). This suggests that signaling filopodia may represent a general mechanism of intercellular communication in living organisms.

Distinguished by their cytoskeletal composition, morphology, and signaling mode, these can be categorized into several types with different terms, including cytonemes, airinemes, tunneling nanotubes, and others (Daly et al., 2022; Eom, 2020; Zhang and Scholpp, 2019). All of them are extended by either signal-sending or signal-receiving cells or both, establishing physical contact with their specific target cells. Signaling molecules, including major morphogens, are moved along the protrusions, or packaged into the vesicles and delivered (Kornberg and Roy, 2014b; Roy et al., 2011; Yamashita et al., 2018).

Airinemes were initially identified in pigment cells in developing zebrafish (Eom et al., 2015). Airinemes are frequently extended by unpigmented yellow pigment cells called xanthoblasts, particularly during metamorphic stages when pigment pattern development takes place (Fig. 1A, B). Unlike fully differentiated xanthophores, xanthoblasts display bleb-like bulged membrane structures at their surface, serving as the origin of airineme-vesicles at the tips of the airinemes (Fig. 2A, yellow arrowheads). The initial step of airineme-mediated signaling involves the interaction of a specific skin-resident macrophage subpopulation, termed metaphocytes, with these blebs (Bowman et al., 2023; Eom and Parichy, 2017). This subset of macrophages an essential role in airineme-mediated intercellular communication, a unique feature not reported in other signaling cellular protrusions that typically involve signal-sending and -receiving cells for their signaling events. Live imaging experiments have shown that macrophages engulf the blebs, pulling them as they migrate, with filaments extending behind the migrating macrophages (Bowman et al., 2023; Eom and Parichy, 2017) (Fig. 2A, white arrowhead). Subsequently, macrophages release the blebs (now referred to as airineme vesicles) onto the surface of target melanophores (Eom, 2020).

**Figure 1.**
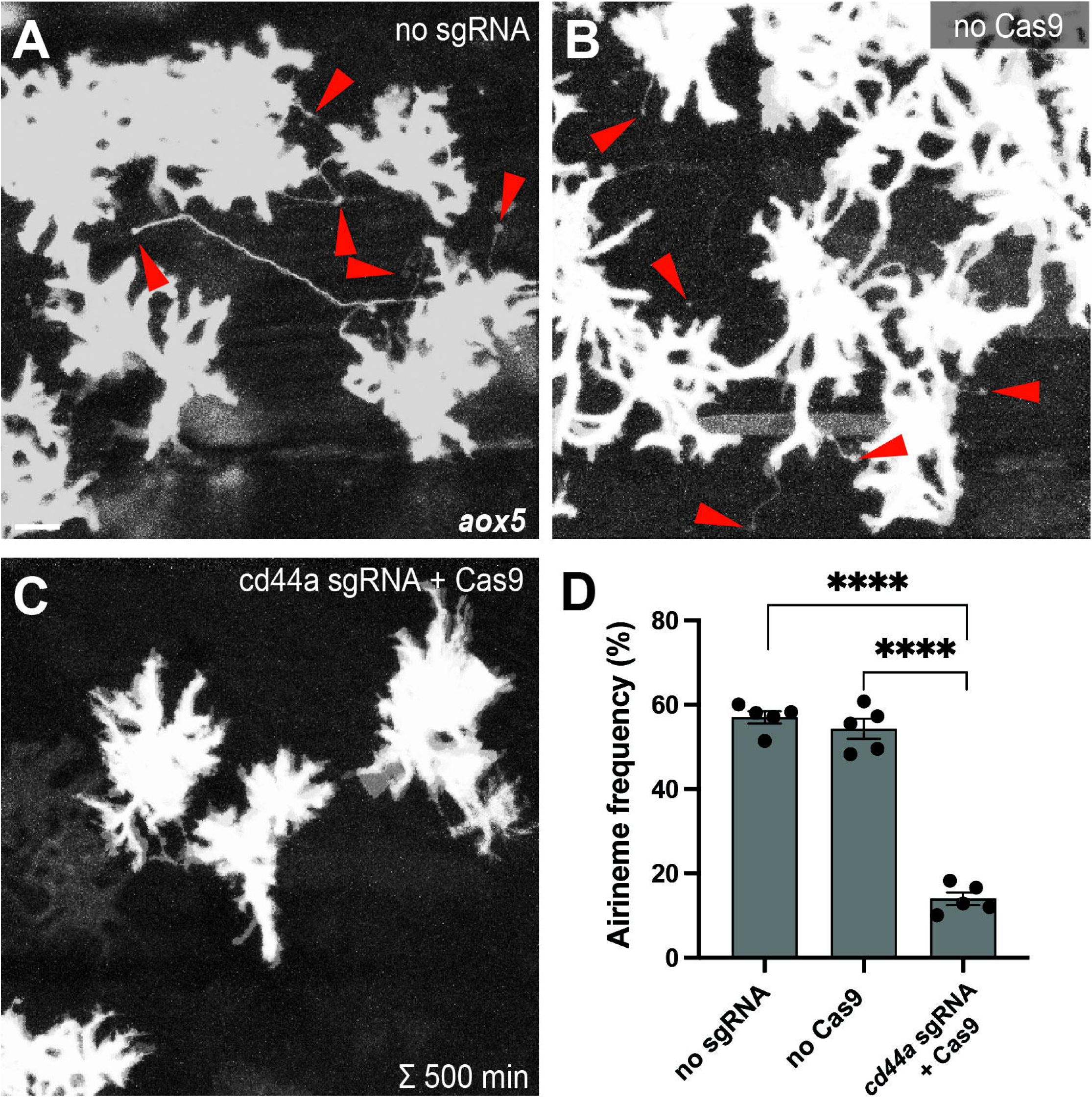
Gene knock-out of cd44a results in significant reduction in airineme extension. (A, B) Merged time-lapse frames over 500 mins display airinemes. Red arrowheads mark airineme vesicles. (C, D) The extension of airinemes was significantly decreased in embryos injected with cd44a sgRNA/Cas9, (*F*_2, 12_) = 172.8, *P*<0.0001, 5 larvae each). Statistically significant results were evaluated using a One-way ANOVA, followed by a Tukey’s HSD post hoc test. Scale bars represent 20μm. Error bars denote mean ± SEM. Images were overexposed to visualize thin airineme filaments.

**Figure 2.**
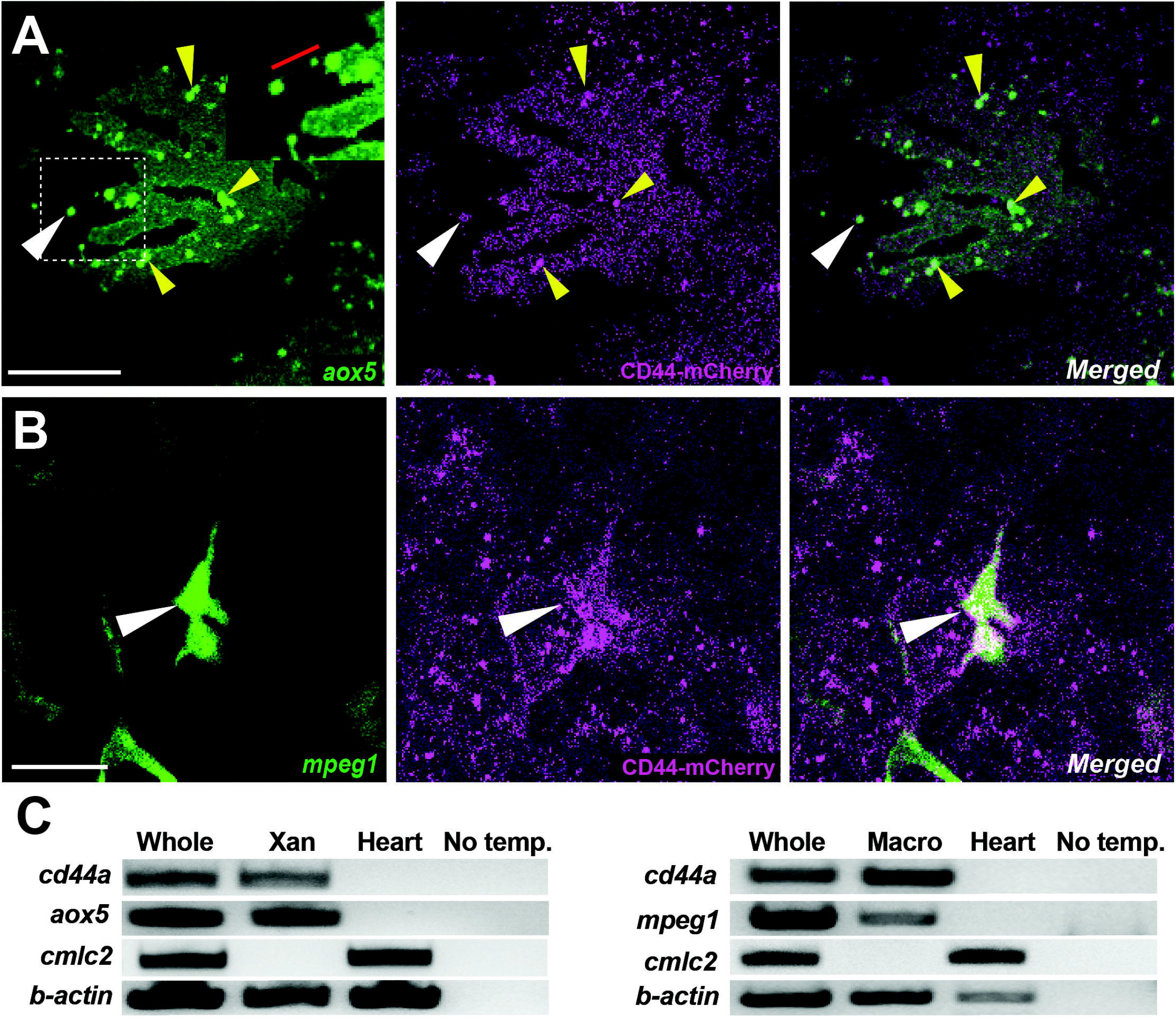
Localization of CD44a protein in xanthoblasts and macrophages. (A) CD44-mCherry expression across the xanthoblasts cell membrane. CD44 is also notably present in the airineme vesicle (white arrowhead) and in the airineme blebs (yellow arrowheads). Airineme filament is indicated with a red bar in the inset. (B) CD44-mCherry expression in macrophages (white arrowheads). (C) RT-PCR for cd44a in isolated xanthophores (xan) and macrophages, and no template control. Whole cDNA was used for positive control. Scale bars represent 20μm (A, B).

The target cells for airinemes are embryonic melanophores and newly differentiating melanophores, rather than fully differentiated melanophores. These target melanophores are situated in the developing interstripe of the metamorphic zebrafish and receive Notch signals through airineme vesicles containing DeltaC ligand. It has been suggested that the Notch signal activated by airinemes subsequently triggers Kita signaling, which is essential for melanophore migration and survival (Eom et al., 2015; Parichy et al., 1999). Consequently, target melanophores within the interstripe gradually migrate out of the interstripe and coalesce into the stripes (Patterson and Parichy, 2019). Inhibiting airineme extension or depleting macrophages leads to the melanophore retention in the interstripe, emphasizing the critical role of airinemes and macrophages in orchestrating pigment patterning during zebrafish development (Bowman et al., 2023; Eom et al., 2015; Eom and Parichy, 2017).

In the initial step of airineme/macrophage-mediated signaling, it is probable that mechanisms exist for macrophages to recognize and adhere to airineme blebs. Previous study showed that airineme blebs express a high level of phosphatidylserine, a phospholipid used as an ‘eat-me’ signal for macrophages to initiate phagocytosis (Davies et al., 2013; Eom and Parichy, 2017; Ginhoux and Guilliams, 2016). However, the necessity and molecular nature of adhesive interactions between these remain unknown.

CD44 is a class I transmembrane adhesion protein with broad expression across various cell types, including lymphocytes, immune cells, fibroblasts, neuronal cells, and more (Ponta et al., 2003). Notably, its overexpression in several cancer stem cells suggests a significant role in cancer development and progression (Chen et al., 2018; Weng et al., 2022). Different CD44 isoforms exhibit distinct functions in interacting with ligands and other interacting molecules, emphasizing their role in diverse tumor progression in humans (Gunthert et al., 1995; Wang et al., 2009).

Like Notch receptors, CD44 can undergo cleavage by membrane type 1 matrix metalloprotease and subsequent cleavage by γ-secretase. This sequential cleavage releases the intracellular domain (ICD) of CD44 into cytosol, allowing its translocation into nucleus. The CD44-ICD functions as a transcription factor that induce genes associated with cell survival, migration, metastasis, and others (Miletti-Gonzalez et al., 2012; Skandalis, 2023). Meanwhile, extracellular domain (ECD) of CD44 facilitates adhesion with other cells expressing CD44 or interact with several ligands in the extracellular matrix (ECM). One of such ligand is hyaluronic acid (HA), and CD44-HA interaction is known to regulate cytoskeleton dynamics and tumor progression in some contexts. Furthermore, CD44-ECD-mediated homophilic cell-to-cell adhesion is well-documented in tumors that can facilitate tumor cell aggregation and metastasis (Kawaguchi et al., 2020; Liu et al., 2019). The multifaceted roles of CD44 highlights its significance in both physiological and pathological cellular processes, making it a crucial focus in cancer research and therapeutic development.

In this study, we demonstrate that CD44, through its extracellular domain, plays a crucial role in the adhesive interaction between the blebs of airineme-producing xanthoblasts and airineme-pulling macrophages. This interaction is critical for airineme-mediated intercellular communication and the formation of pigment pattern in zebrafish.

## Results

### Gene knock-out of cd44a using CRISPR/Cas9 results in a substantial decrease in airineme extension

To identify candidate genes involved in adhesive interaction between macrophages and airineme vesicles, we conducted gene expression profiling between xanthophores and airineme-producing xanthoblasts. Among several adhesion proteins analyzed, cd44a exhibited the most significant gene expression difference between these two cell types (log2 fold change of 10.13). To investigate the role of cd44a in airineme-mediated signaling, we designed single-guide RNA (sgRNA) against cd44a and injected it into one-cell-stage embryos with Cas9 protein, along with an *aox5:palmEGFP* construct to label cell membrane and airinemes of xanthophore-lineages. Control groups included embryos that receive cd44a sgRNA injection without Cas9 protein and embryos injected only with Cas9 protein into wild-type embryos. The fish were raised until metamorphic stages (SSL 7.5), when airineme extension is most frequent (Eom et al., 2015; Parichy et al., 2009). We counted the number of cells extending airinemes out of the total cells imaged at 5-minute intervals over a period of 10 hours during overnight time-lapse imaging (Eom et al., 2015). This method was also used for subsequent quantifications to measure airineme extension frequency. Our results showed that embryos injected with cd44a sgRNA/Cas9 had a significant reduction in airineme extension as compared to the two controls, suggesting that cd44a may be required for proper airineme signaling (Fig. 1).

### CD44 expression in macrophages and xanthophore-lineages

The role of CD44 in cell-cell or cell-ECM interactions is well-established in various contexts (Chen et al., 2018; Ponta et al., 2003; Senbanjo and Chellaiah, 2017). Given this, we predicted that CD44 is expressed in either airineme producing xanthoblasts or macrophages, or potentially both. To investigate the localization of CD44 protein under native regulatory elements, we recombineered an 82 kb BAC (Bacterial Artificial Chromosome) containing zebrafish cd44a coding sequence and regulatory elements to generate an mCherry fusion, *TgBAC(cd44a:cd44a-mCherry)*. To assess CD44 protein expression in xanthoblasts, we injected *aox5:palmEGFP* construct into *TgBAC(cd44a:cd44a-mCherry)*. The CD44-mCherry signal was detected in various cell types, including xanthophore lineages (Fig. 2A). Interestingly, we were able to detect enriched CD44 expression in the airineme vesicles (Fig. 2A, white arrowhead) and airineme blebs (Fig. 2A, yellow arrowheads, Fig S1) which are the precursors of airineme vesicles. To examine CD44 expression in macrophages, we injected *mpeg1:palmEGFP* construct, which label cell membranes of all macrophages. We confirmed that macrophages, specifically metaphocytes, also express CD44 (Fig. 2B, arrowheads) (Vachon et al., 2006). Metaphocytes were easily distinguishable by their amoeboid morphology compared to the dendritic population in the zebrafish skin (Bowman et al., 2023; Kuil et al., 2020; Lin et al., 2020; Lin et al., 2019). CD44-mCherry expression was not confined to specific subcellular structures except airineme blebs but was distributed throughout the cells in xanthophore-lineages. Given the fact that CD44 is a membrane protein, we assumed its expression at least in the cell membrane.

Although, macrophages exhibited strong CD44 expression at the center of the cell, it was also expressed throughout the cells. Thus, our findings confirm that CD44 protein is expressed in both xanthoblasts and macrophages (Fig. 2A, B). We also collected EGFP-labelled xanthophore lineages from *Tg(aox5:palmEGFP)* and macrophages from *Tg(mpeg1:palmEGFP)* using Fluorescence-activated Cell Sorting (FACS). We confirmed cd44 mRNA expression in both cell types using RT-PCR. For negative controls in this experiment, we tested multiple tissues and organs in zebrafish. We found that heart tissue consistently lacked cd44 expression and thus used it as a negative control (Fig. 2C).

### Extracellular domain of CD44 is essential for airineme extension

As CD44 is a multifunctional protein that can promote adhesion between cells or extracellular matrix (ECM) through interactions with various ligands one well-known ligand for CD44 is hyaluronic acid (HA). (Senbanjo and Chellaiah, 2017). Given the fact that airinemes are actin- and tubulin-based structures, and the CD44-HA interaction can activate the cytoskeleton in certain contexts, we aimed to investigate whether CD44 controls airineme extension through its interaction with HA (Eom, 2020; Eom et al., 2015; Ponta et al., 2003). We anticipated observing HA expression in CD44-expressing xanthoblasts if CD44 acts through HA. However, we did not detect any overlapping expression of HA in xanthoblasts, suggesting that the CD44-HA interaction does not appear to be involved in airineme signaling (Fig. S2).

Domain studies of CD44 suggested that its intracellular domain (ICD) functions as a transcription factor, influencing cell migration, angiogenesis, invasion and other behaviors (Chen et al., 2018; Skandalis, 2023). To investigate whether CD44-ICD plays a role in airineme extension, we generated transgenic lines overexpressing cd44-ICD in either xanthophore-lineages or macrophages, *Tg(aox5:cd44aICD)* and *Tg(mpeg1:cd44aICD)*. In both transgenic lines, we did not observe any significant changes in airineme extension frequency, suggesting that CD44-ICD doesn’t seem to be involved in airineme extension (Fig. 3A, B-B”, E).

**Figure 3.**
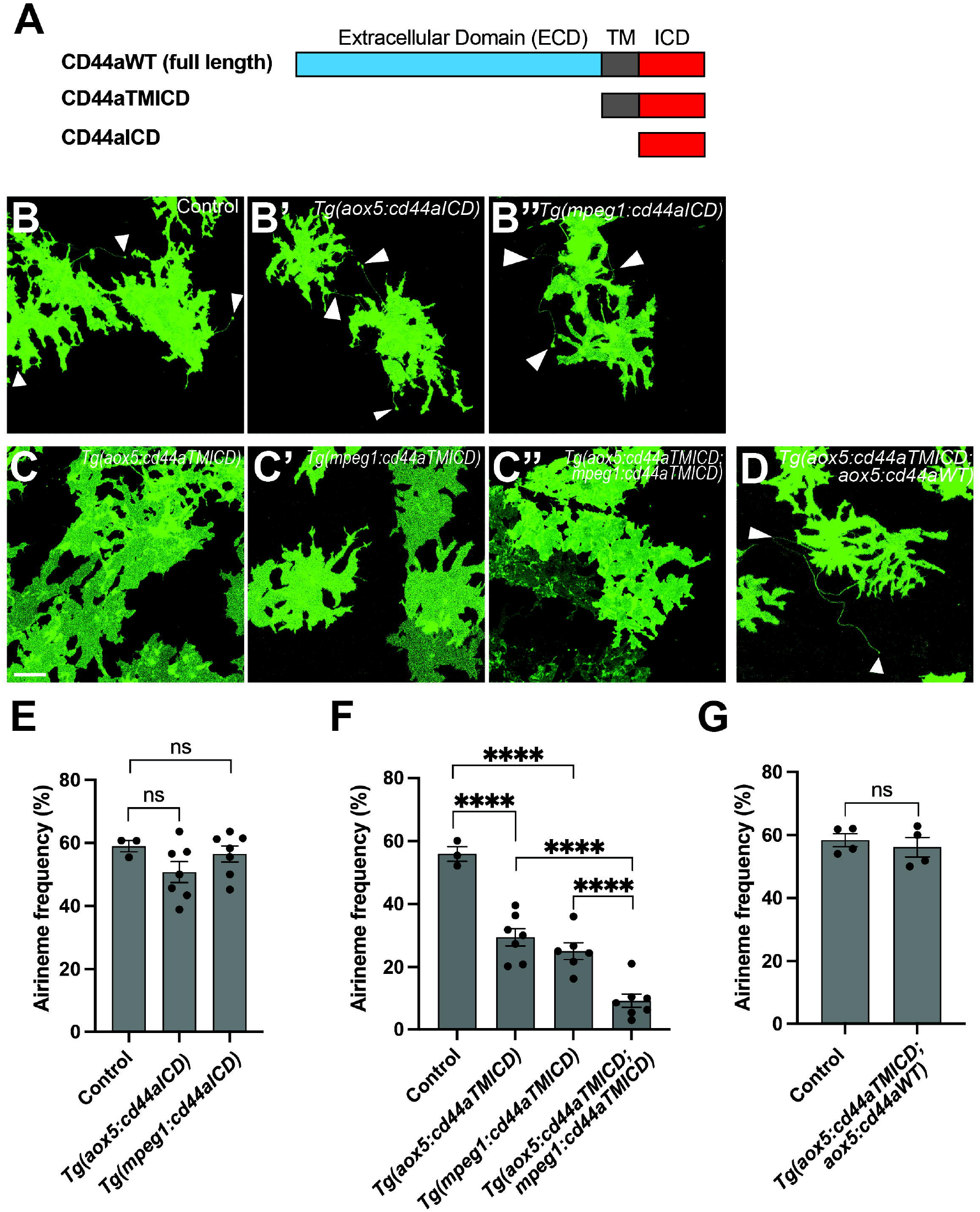
The crucial role of the extracellular domain of CD44 in airineme extension. (A) The schematics indicating the structure of CD44aWT, CD44aTMICD, and CD44aICD. (B-B”) Representative images of xanthophores extending airinemes in control, *Tg(aox5:cd44aICD)*, or *Tg(mpeg1:cd44aICD)*. (C-C”) Representative images of xanthophores in *Tg(aox5:cd44aTMICD), Tg(mpeg1:cd44aTMICD)*, or *Tg(aox5:cd44aTMICD; mpeg1:cd44aTMICD)*. (D) Representative image of xanthophores in *Tg(aox5:cd44aTMICD; aox5:cd44aWT)*. White arrowheads mark airinemes (E) Airineme extension frequency did not significantly altered in transgenic embryos that overexpressed the intracellular domain of cd44 (cd44aICD) specifically in the xanthophore-lineages or macrophages, (*F*(_2, 14_)=1.736, *P*=0.2121, 17 embryos in total). (F) Overexpression of CD44a with a truncated extracellular domain (cd44aTMICD) exhibited significant reduction in airineme extension frequency in both the xanthophore-lineages and macrophages, (*F*(_2, 13_) = 23.62, *P*<0.0001, 16 embryos in total). Airineme extension frequency was even further significantly reduced when cd44aTMICD was overexpression in both cell type simultaneously, (*F*(_2, 17_)=18.04, *P*<0.0001). (G) Airineme extension frequency was restored when the WT CD44 was overexpressed in embryos already expressing cd44aTMICD in the xanthophore lineages, (*P*=0.5767, 4 embryos each). Statistically significant results were evaluated using a One-way ANOVA, followed by a Tukey’s HSD post hoc test or a Student’s t test. Error bars indicate mean ± SEM.

Subsequently, we explored the potential role of the extracellular domain (ECD) of CD44 in airineme extension. CD44-ECD is known to interact with the extracellular matrix (ECM) or CD44-ECD from other cells (Senbanjo and Chellaiah, 2017). To test this, we generated transgenic lines overexpressing an ECD-truncated form of cd44 (cd44TMICD), specifically in xanthophore-lineages, *Tg(aox5:cd44aTMICD)* or macrophages, *Tg(mpeg1:cd44aTMICD)*. These transgenic lines did not impact cell viability or other noticeable cellular behaviors in either xanthophore-lineages or macrophages (Fig. S3). Interestingly, we observed a significant decrease in airineme extension frequency in the transgenic lines overexpressing cd44aTMICD (Fig. 3A, C-C”, F). We investigated whether the additional supply of WT CD44 could rescue the phenotype in *Tg(aox5:cd44aTMICD)* by generating a double transgenic line, *Tg(aox5:cd44aTMICD; aox5:cd44aWT)*. These rescue fish exhibited almost full recovery of airineme extension frequency. Together, these results suggest that CD44-ECD is critical for airineme extension (Fig. 3A, D, G).

### Trans-adhesive interaction via CD44ECD between xanthoblasts and macrophages is critical in airineme extension

We investigated whether extracellular domain of zebrafish CD44 can mediate adhesion between cells. To test this, we expressed constructs that encode wild-type zebrafish cd44a or ECD-truncated cd44a (cd44aTMICD), both C-terminally fused with mCherry, in *Drosophila* S2 cells (Schneider, 1972). Actin-mCherry fusion construct was used as a control. Transfected S2 cells expressing mCherry were monitored, and after several hours of culture in a rotary incubator, we assessed whether CD44 expressing S2 cells adhere to each other. The results showed that, unlike the actin control and cd44aTMICD-expressing S2 cells, wild-type cd44a-expressing S2 cells began forming aggregates 30 minutes after incubation. The degree of aggregation increased over time, reaching a significant level at 3 hours. This indicates that zebrafish CD44 mediates trans-adhesion via its ECD between cells (Fig. 4A, B).

**Figure 4.**
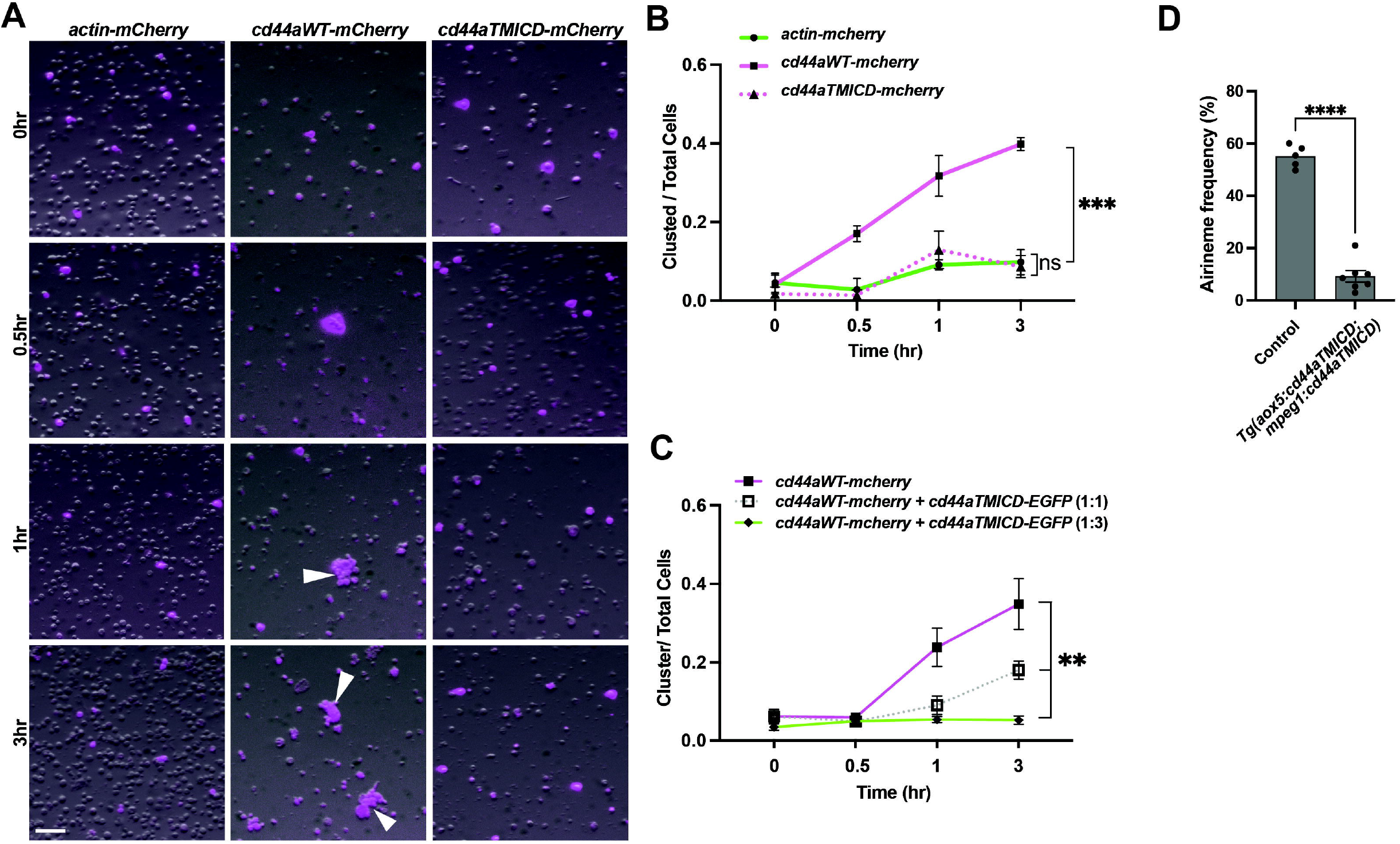
Trans-adhesive interaction via extracellular domain of CD44. (A) S2 cells were transfected with constructs to express actin-mCherry (control), wild-type cd44a-mCherry, or cd44aTMICD-mCherry. Within 1 hour of rotary culture, cell aggregates were formed by those expressing the wild-type cd44a (white arrowheads). (B) Quantification of the transfected cells found within clusters relative to the total numbers of transfected cells in 3 replicate cultures, (*F*(_2, 6_)=45.50, *P*=0.0002). (C) S2 cells were co-transfected with constructs expressing wild-type cd44a-mCherry and cd44aTMICD-EGFP at ratios of 1:0 (control), 1:1 and 1:3, (*F*(_2, 6_)=13.7, *P*=0.0058). Corresponding images can be found in Fig. S3. (D) Overexpressing cd44aTMICD in both xanthophore lineages and macrophages resulted in a significant reduction in airineme extension frequency. Statistically significant results were evaluated using one-way ANOVA, followed by a Tukey’s HSD post hoc test or a Student’s t-test. Scale bars: 100μm. Error bars indicate mean ± SEM.

Previously, we demonstrated that overexpressed cd44aTMICD inhibits airineme extension frequency. This effect was rescued by the supply of WT cd44a *in vivo* (Fig. 3B, C). To further investigate whether the overexpression of CD44aTMICD in the presence of WT CD44a could negatively affect cell adhesion, we co-expressed *cd44aWT-mCheery* and *cd44aTMICD-EGFP* in S2 cells. We observed that the degree of S2 cell aggregation significantly decreased as the proportion of CD44aTMICD-EGFP expression relative to CD44aWT-mCherry increased (Fig. 4C, S4). Taken together with the *in vivo* data in Figure 3, these results suggest that overexpressed CD44aTMICD exhibited a dominant negative effect by interfering with the ability of wild-type CD44a to mediate cell adhesion. CD44TMICD might disrupt the organization of membrane microdomains, protein complexes, or dimers / oligomers necessary for CD44 function (Kawaguchi et al., 2020; Ma et al., 2022). However, the underlying mechanism of this interference requires further investigation.

These findings prompted us to ask whether trans-adhesive interactions between xanthoblasts and macrophages could play a role in airineme extension in zebrafish. We hypothesized that a greater reduction in airineme extension frequency would be observed if CD44a function is compromised simultaneously in both cell types, compared to its expression in only one cell type. To test this, we generated a double transgenic line that overexpresses cd44aTMICD in both xanthophore-lineages and macrophages, *Tg(aox5:cd44aTMICD; mpeg1:cd44aTMICD)*. Indeed, we observed a more significant decrease in airineme extension in this double transgenic zebrafish compared to single manipulations (Fig. 3B, 4D), suggesting, at least in part, a requirement for CD44ECD by both xanthoblasts and macrophages for proper airineme signaling as CD44 is expressed in airineme vesicles and macrophages (Fig. 2A, B). However, we cannot rule out the possibility that there are other adhesion molecules may trans-interact with CD44 in both xanthophore lineages and macrophages.

Next, we asked whether CD44-mediated adhesion is critical when macrophages contact airineme blebs (equivalent to airineme vesicle) and or when they drag the airineme vesicles. We hypothesized that if such adhesive interaction is important to maintain the attachment of airineme vesicles to macrophage while dragging, a reduction in airineme length would be observed. This is because the detachment of macrophages from the airineme vesicle would stop the extension of airineme filaments (Bowman et al., 2023; Eom and Parichy, 2017). However, we did not detect any statistically significant change in airineme length in embryos overexpressing CD44aTMICD in either cell type or both simultaneously (Fig. S5). This finding suggests that the trans-adhesive interaction mediated by CD44 plays a more significant role when macrophages are contacting airineme blebs as opposed to pulling the airineme vesicles.

### CD44-mediated airineme signaling is crucial for pigment pattern formation

Airineme-mediated intercellular signaling is indispensable for pigment pattern formation in zebrafish. Our results indicating the importance of CD44a in airineme extension led us to anticipate pigment pattern defects in embryos overexpressing cd44aTMICD. However, noticeable pigment pattern defects were not detected in either transgenic line overexpressing cd44aTMICD in xanthophore lineages or macrophages. This suggests that residual airineme extensions may still be sufficient to deliver the necessary signals for generating a stripe pattern, or subtle difference could be challenging to detect (Fig. 2B). Nevertheless, when CD44a function was compromised by overexpressing cd44aTMICD in both cell types, a significant number of melanophores were observed to be retained in the interstripe compared to the control, although the total number of melanophores remained unchanged (Fig. 5A, B, C). We also observed the consistent pigment pattern defects from cd44 mutants induced by CRISPR/Cas9 system (Fig. S6).

**Figure 5.**
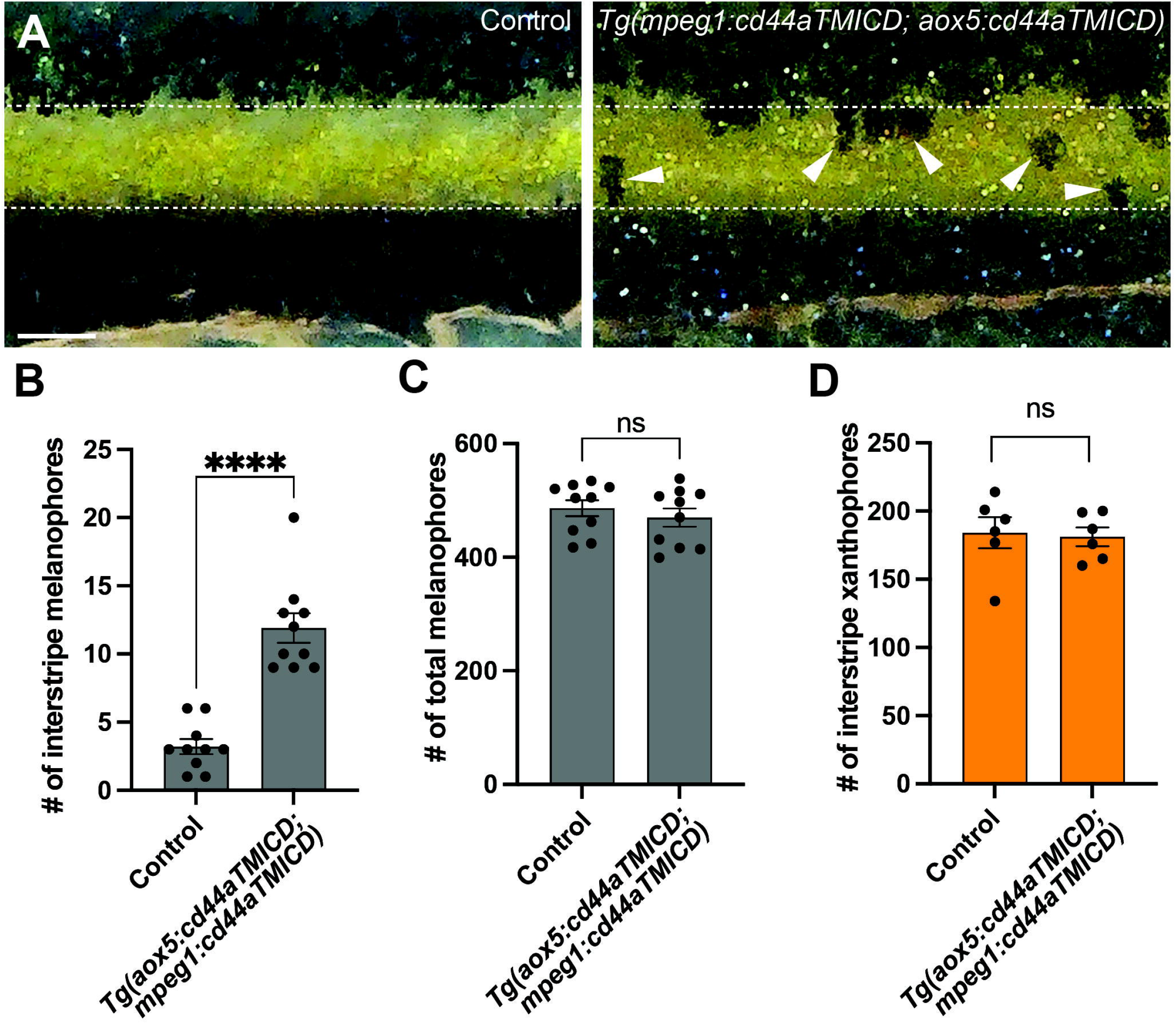
CD44-mediated airineme extension contributes to zebrafish pigment pattern formation. (A) Unlike the control, melanophores failed to coalesce into stripe and remained in the interstripe zone (white arrowheads) at SSL11 (Parichy et al., 2009). The white dotted lines demarcate stripes and interstripe. (B) In embryos overexpressing cd44aTMICD simultaneously in both xanthophore-lineages and macrophage, the count of interstripe melanophores was significantly higher, (*P*<0.0001, 20 embryos in total). (C) However, the total number of melanophores did not differ significantly between the experimental group and controls, (*P*=0.4525, 20 embryos in total). (D) The number of xanthophores in the interstripe did not differ significantly between the experimental group and controls (*P*=0.8266, 12 embryos total). Statistical significance was assessed using a Student’s t test. Scale bars represent 200μm. Error bars indicate mean ± SEM.

Together, these findings collectively suggest that CD44-mediated trans-adhesive interaction between airineme blebs on xanthoblasts and macrophages play an essential role in airineme-mediated signaling during pigment pattern formation in zebrafish.

## Discussion

In this study, we demonstrated that CD44 protein is expressed in xanthoblasts, particularly in the airineme vesicles (Fig. 2A, yellow arrowheads). Intriguingly, though, not every airineme bleb expresses CD44 but only some of them do. This observation leads to a few possible explanations. One explanation could be that CD44 expression determines which blebs are used for airineme signaling. It is possible that some blebs do not express CD44 and serve a different function beyond airineme signaling. In addition, our observation may suggest that there is a mechanism by which CD44 protein in the cell membrane becomes further enriched in the airineme blebs. Similar to CD44, we observed that DeltaC, one of signaling molecule found in the airineme vesicles, is not expressed in some of the blebs before their extraction by metaphocytes (Eom et al., 2015). It is conceivable that there are mechanisms that sort proteins required for airineme signaling into the blebs and ultimately into the airineme vesicle. These might be used to adhere to the macrophage membrane and activate Notch signal on the target cell surface. It would be interesting to study the underlying molecular mechanisms involved in sorting out those proteins essential for airineme-mediated intercellular signaling into the airineme blebs.

Furthermore, CD44 has previously been reported as a signal for phagocytosis in macrophages (Vachon et al., 2007; Vachon et al., 2006). Therefore, alongside phosphatidylserine (PtdSer), CD44 may serve as an additional recognition signal for macrophages to identify airineme blebs, as well as acting as an adhesive signal between airineme blebs and macrophages (Eom and Parichy, 2017).

This raises an interesting question as to why we did not observe any differences in airineme filament length when the adhesive interaction mediated by CD44 was reduced (Fig. S5A). Our interpretation is that this adhesive interaction would be crucial during the initial interaction between macrophages and the airineme blebs, rather than during the process of dragging them. Thus, it seems plausible that CD44 functions as both a recognition and adhesion element at the time when macrophages interact with airineme blebs. On a related note, our previous research, using high-resolution 3D confocal image reconstruction, has shown that airineme vesicles are found inside the dragging macrophages (Eom and Parichy, 2017). This implies that airineme vesicles are physically entrapped within the macrophages during the pulling process, suggesting that adhesive interaction between the membranes of those two signaling components might not be strictly necessary while pulling. Thus, it is conceivable that CD44 might play a dual role. First, it may assist PtdSer in recognizing airineme blebs and simultaneously provide necessary adhesion during the recognition and subsequent engulfment by the macrophage but not necessary while airineme vesicles are dragged by macrophages.

CD44 could function as a member of a functional membrane microdomain facilitating this interaction. However, future experiments would be required to verify this hypothesis.

We observed residual airineme extension even when we overexpressed cd44aTMICD in both xanthophore lineages and macrophages (Fig. 4C). This residual airineme extension may indicate that either remaining wild-type CD44 continues to mediate adhesion or other adhesion proteins and players are involved in the process. To explore this further, we considered the studied interaction of Matrix Metalloproteinase-9 (MMP9) with CD44 (Senbanjo and Chellaiah, 2017). Cell surface expression of MMP9, along with CD44, is known to facilitate cell migration and invasion in tumors and PC3 cells (Desai et al., 2008; Gupta et al., 2013; Yu and Stamenkovic, 1999). Given our previous findings that airineme-pulling macrophages express high-levels of mmp9, crucial for macrophage migration and penetration into the hypodermis in zebrafish skin (Bowman et al., 2023), we explored whether CD44 also plays a role in macrophage migration speed (=airineme extension speed) through its interaction with MMP9. However, our analysis did not reveal a significant difference in airineme extension speed in cd44aTMICD overexpressed macrophages (Fig. S5B). Although further investigations are necessary, this suggests that CD44 may function independently in macrophage migration during airineme extension.

Taken together, our study discovered that extracellular domain of CD44 in both airineme-extending xanthoblasts and airineme-pulling macrophages facilitates a trans-adhesive interaction, which appears to be critical in initiating airineme extension. This study provides evidence for the requirement of cellular adhesion between signaling-sending cell and relaying cells in airineme-mediated intercellular signaling. It is also conceivable that similar mechanisms could be conserved in other cellular contexts.

## Materials Availability

This study generated several zebrafish transgenic lines, *Tg(mpeg1:cd44aICD-v2a-mCherry), Tg(aox5:cd44aICD-v2a-mCherry), Tg(mpeg1:cd44aTMICD-v2a-mCherry), Tg(aox5:cd44aTMICD-v2a-mCherry), Tg(aox5:cd44aFL-v2a-mCherry), Tg(mpeg1:cd44aFL-v2a-mCherry)* and *TgBAC(cd44a:cd44amCherry)*. They are available from the lead contact without restriction.

### Experimental Model and Subject Details

#### Zebrafish

Fish were maintained at 28.5C, 16:8 L:D. Zebrafish were wild-type AB^WP^ or its derivative WT(ABb), as well as *Tg(mpeg1:Brainbow*^*W201*^), which expresses tdTomato in the absence of Cre-mediated recombination, and *Tg(aox5:palmEGFP)*. Experiments were performed prior to the development of secondary sexual characteristics, so the number of males and females in the study could not be determined, however, all stocks generated approximately balanced sex ratios, so the experiments likely sampled similar numbers of males and females. All animal work in this study was conducted with the approval of the University of California Irvine Institutional Animal Care and Use Committee (Protocol #AUP-25-002) in accordance with institutional and federal guidelines for the ethical use of animals.

## Methods

### Transgenesis and transgenic line production

To examine how loss of extracellular domain (ECD) of CD44 adhesion molecule may impact interactions between macrophages and xanthoblasts, we generated *mpeg1:cd44aTMICD-v2a-mCherry* and *aox5:CD44aTMICD-v2a-mCherry* constructs using Gateway assembly into Tol2 backbone and injected into WT(ABb). Truncated version of CD44a (CD44TMICD) was obtained by extracting CD44aTMICD cDNA from WT(ABb) cDNA. To examine whether interactions between macrophages and xanthoblasts were gene expression dependent via CD44a-ICD, we generated *Tg(aox5:CD44aICD-v2a-mCherry)*^*ir*.*rt8*^ and Tg(*mpeg1:CD44aICD-v2a-mCherry)*^*ir*.*rt10*^ transgenic lines, which were likewise generated using Gateway assembly into Tol2 backbone and injected into WT(ABb). Similarly, CD44aICD cDNA was isolated from WT(ABb) cDNA. To further examine how ECD of CD44a may impact interactions between macrophages and xanthoblasts, we generated *Tg(aox5:CD44aFL-v2a-mCherry)*^*ir*.*rt22*^ to see if we could rescue the phenotypes seen *in Tg(aox5:CD44aTMICD-v2a-mCherry)*^*ir*.*rt9*^.

To visualize CD44 localization under native regulatory elements, we inserted mCherry C-terminally using BAC CH211-102L7 with 82 kb 5’ and 92 kb 3’ to the open reading frame.

### Hyaluronic Acid Binding Protein (HABP) assay

Zebrafish embryos (SSL7.5) were incubated in a fish water diluted biotin-HABP (1:150, amsbio) along with streptavidin-Alexa546 for 1 hr. Controls incubated solely with streptavidin-Alexa546. Fish were then washed two times with fish water and imaged under a confocal microscope.

### Time-lapse imaging and still imaging

*Ex-vivo* imaging of pigment cells and macrophages was done using Leica TCS SP8 confocal microscope equipped with a resonant scanner and two HyD detectors. Time-lapse images were taken at 5-min intervals over a span of 10h. Overnight time-lapse imaging was performed when larvae reached SSL (Standardized Standard Length) 7.5 (Parichy et al., 2009).

### S2 Cell Adhesion Assay

S2 cells were obtained from the Drosophila Genomics Resource Center (DGRC, S2-DGRC (DGRC Stock 6 ; https://dgrc.bio.indiana.edu//stock/6 ; RRID:CVCL_TZ72)) and cultured in Schneider’s Drosophila Medium (Catalog No. 21720024, Gibco, USA) supplemented with 10% Fetal Bovine Serum (FBS) (Catalog No. 25-514H, Genclone, USA) and 1% Penicillin-Streptomycin (Catalog No. 15140148, Gibco, USA). For transfection, S2 cells were transfected with three different plasmids: *pAW-actin-mcherry, pAW-cd44a-FL(wild type full length)-mcherry*, and *pAW-cd44a-TMICD-mcherry*. Lipofectamine™ LTX Reagent (Catalog No. 15338030, Invitrogen, USA) was used for transfection following the manufacturer’s instructions. After 72 hours of transfection, the cells were counted, and each type of cell was seeded into 35mm glass-bottom petri dishes at a density of 5000 cells per dish (Catalog No. 706011, Nest Scientific, USA). The dishes were then subjected to rotating incubation for different time intervals, including 30 minutes, 1 hour, and 3 hours. Cellular imaging was performed using a Zeiss microscope equipped with a 599nm filter to capture the cellular mCherry signal and bright field images.

To evaluate the outcompeting ability of cd44a-TMICD against the wild type cd44a, S2 cells were co-transfected with *pAW-cd44a-FL-mcherry* and *pAW-cd44a-TMICD-EGFP* at ratios 1:1 and 1:3, using the same transfection protocol described above. After 72 hours of transfection, 5000 cells were transferred into 35mm glass-bottom petri dishes and subjected to rotating incubation at intervals of 30 minutes, 1 hour, and 3 hours.

Cellular imaging was then conducted using a Zeiss microscope under 488nm or 599nm filters to capture the EGFP and mCherry signals, along with bright field images.

### Reverse Transcription Polymerase Chain Reaction (RT-PCR) Analysis of Gene Expression in Zebrafish Skin

Metamorphic stage fish were skinned (n = 20 per group) and subsequently washed in 1x PBS solution. After washing, the samples were briefly centrifuged, and the supernatant was discarded by pipetting. The tissues were resuspended in 1mL of Stem® Pro® Accutase Cell Dissociation Reagent (Gibco) and incubated for 10 minutes at 37°C, or until the cells were dissociated entirely. Following incubation, the cells were passed through a 40μm cell strainer to remove non-dissociated tissue before running through FACS (Fluorescence-Activated Cell Sorting). Using FACS, cells of interest were separated and collected based on their respective membrane markers. Cells containing membrane markers for EGFP were collected for downstream non-quantitative RT-PCR. Following collection, cDNA was synthesized using the SuperScript III CellsDirect™ cDNA synthesis kit (Invitrogen). Non-quantitative RT-PCR amplifications were performed with 40 cycles (*actb1, aox5, mpeg1, cd44a, cmlc2*), utilizing PrimeStar GXL DNA Polymerase (Takara).

*actb1*: 5’-CATCCGTAAGGACCTGTATGCCAAC-3’, 5’-AGGTTGGTCGTTCGTTTGAATCTC-3’, *aox5*: 5’-AGGGCATTGGAGAACCCCCAGT-3’, 5’-ACACGTTGATGGCCCACGGT-3’, *mpeg1*: 5’-CCCAGTGTCAGACCACAGAAGATGGAGTC-3’, 5’-CATCAACACTTGTGATGACATGGGTGCCG-3’, *cd44a*: 5’-GCTGTACTTCAGGCAGCCCC-3’, 5’-GTTTGGACCATTAATGTGTGGGAGG-3’, *cmlc2*: 5’-CAAGAGGGGGAAAACTGCTCAAAG-3’, 5’-GCAGCAAGGATGGTTTCCTCTG-3’,

*mCherry*: 5’-ATGGTGAGCAAGGGCGA-3’, 5’-TTACTTGTACAGCTCGTCCATG-3’

### Generation of *cd44a* mutants by the CRISPR/Cas9 system

Mutations of *cd44a* (accession number XM_001922456) were induced by CRISPR/Cas9. Using SMART, a target site was generated in exon 2, with the following sequence 5’ - GGTGAACTGTGTCAGAGTTT - 3’. Single guide RNA (sgRNA) was then synthesized using MEGAshortscriptTM T7 High Yield Transcription Kit (Invitrogen) and co-injected into one-cell stage embryos with 1μg/μL TrueCutTM Cas9 Protein v2 (InvitrogenTM) at a concentration of 300ng/uL.

### Pigment pattern and melanophore counts

Fish were imaged upon reaching SSL 11.0 using a Koolertron LCD digital microscope. Images were taken of the entire trunk and were later cropped to include only the area underneath the dorsal fin. ImageJ cell counter plugin was used to count interstripe and total numbers of melanophores. Numbers for each group were averaged.

### Quantification and statistical analysis

Statistical analyses were performed using GraphPad Prism.

## Acknowledgements

We thank Parichy lab for their support in the preliminary stages of this study. Additionally, we thank Zach Waller for his contribution to generating and managing some of the transgenic lines used for this study. We acknowledge funding from NIH R35GM142791 to D.S.E. and support from the Drosophila Genomics Resource Center (NIH Grant 2P40OD010949).

