## Supplementary Figures for "CD44 facilitates adhesive interactions in airineme-mediated intercellular signaling"

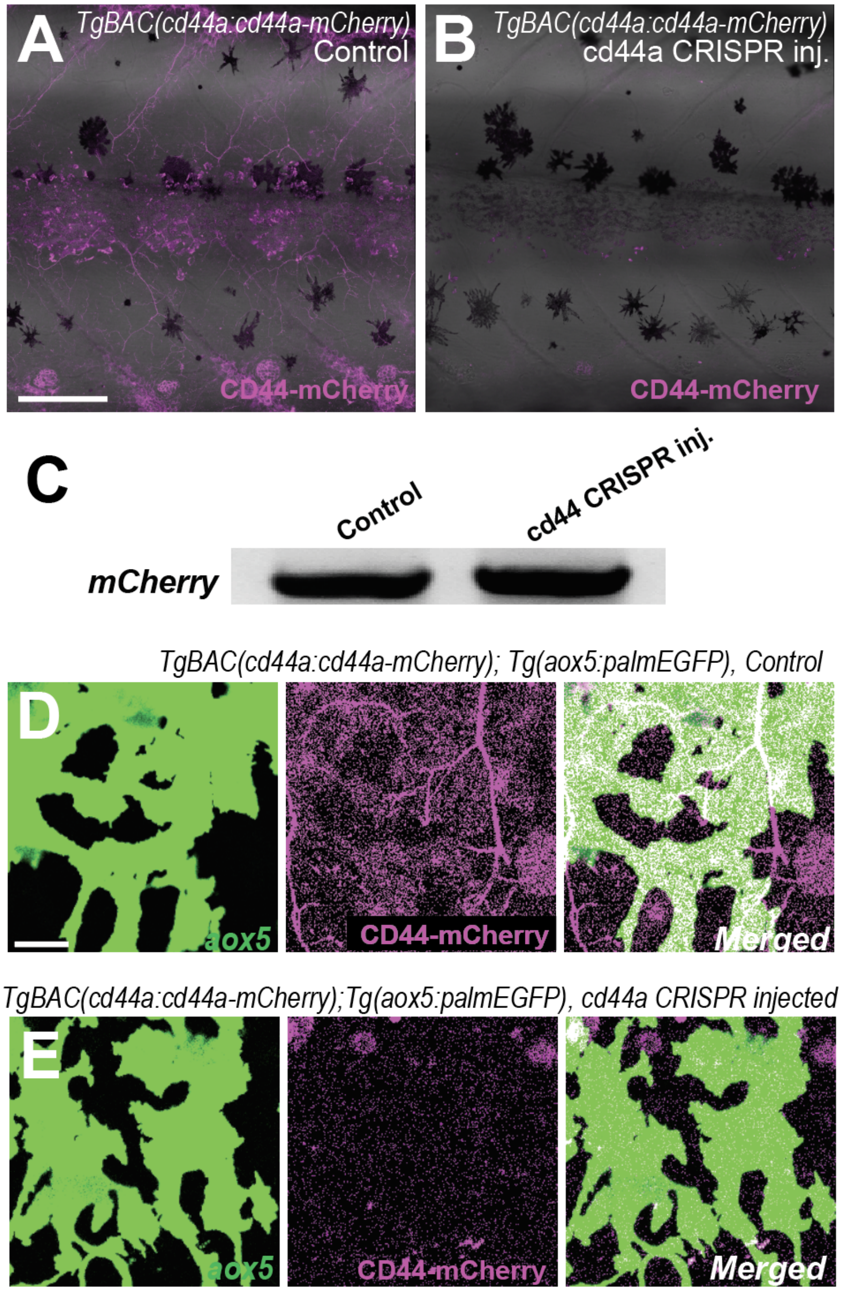


**Figure S1. Verification of CD44-mCherry signal**
To confirm that the CD44-mCherry expression observed in *TgBAC(cd44:cd44-mCherry)* is not a background signal, we injected cd44 CRISPR/Cas9 into *TgBAC(cd44:cd44-mCherry)* fish. (A) Expression of CD44-mCherry was not altered in Cas9-only control fish. (B) A dramatic reduction or undetectable CD44-mCherry signal was observed in cd44 CRISPR/Cas9 injected fish. Note that some autofluorescent signals are present. (C) Both control and injected fish were verified as *TgBAC(cd44:cd44-mCherry)* by detecting mCherry expression. (D) High magnification images with xanthophore marker (aox5) showed CD44-mCherry expression in xanthophores in control. (E) CD44-mCherry signal was not visible in the cd44a CRISPR/Cas9-injected fish. Scale bar represents 200μm (A, B) and 20μm (D, E).

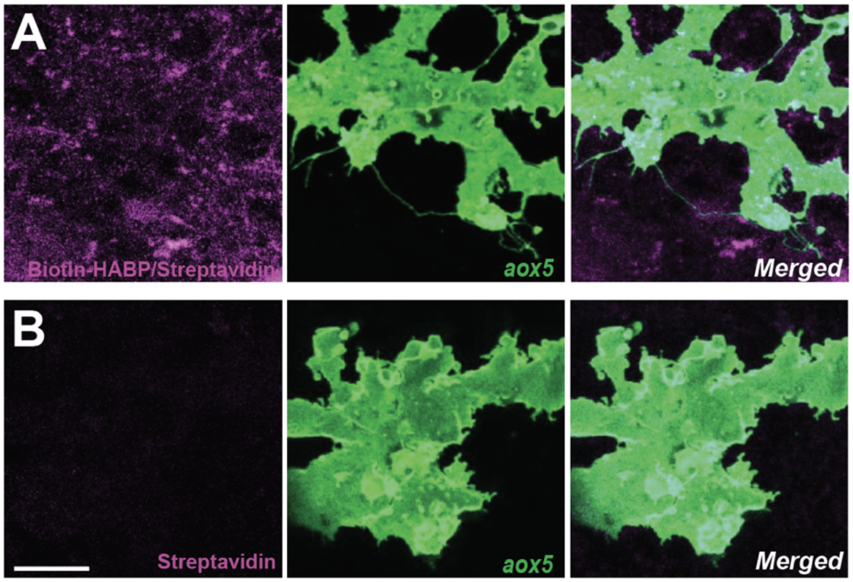


**Figure S2. Lack of apparent Hyaluronic acid binding protein (HABP) expression in xanthoblasts**
(A) Xanthoblasts appear to show no overlapping expression when traced using Biotin-conjugated HABP and Alexa-546-labelled streptavidin. (B) A control experiment with Alexa-546 streptavidin alone did not result in any detectable signal. Scale bar represents 20µm. Images were overexposed to visualize thin airineme filaments.

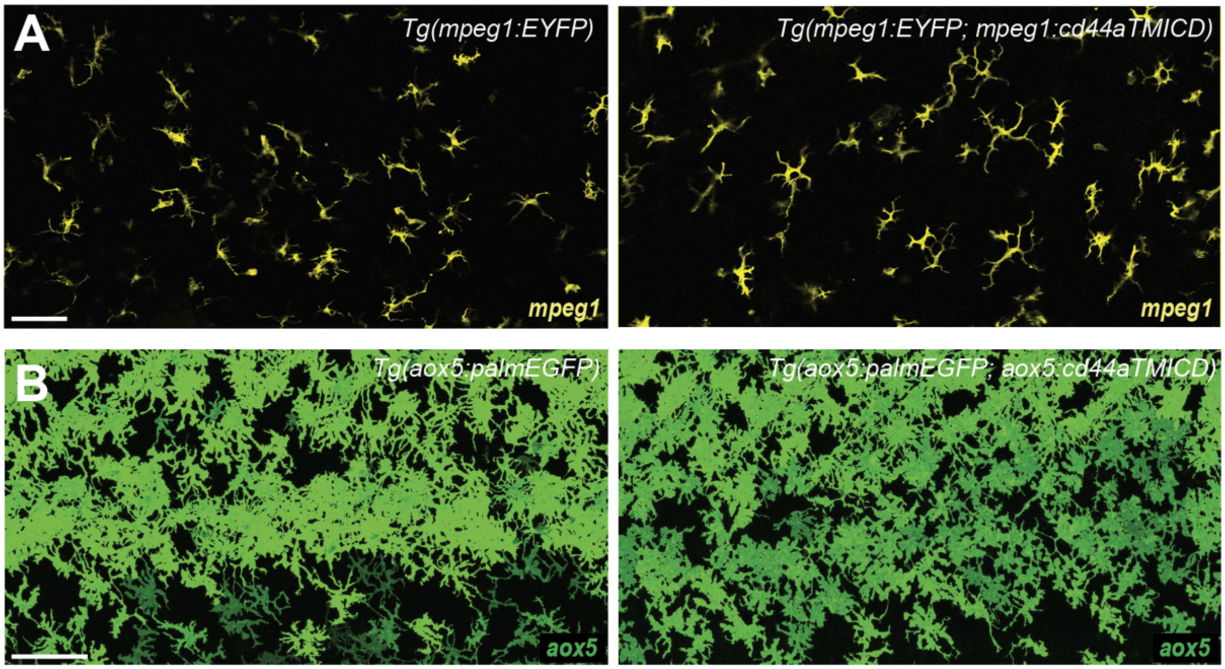


**Figure S3. Overexpression of cd44aTMICD does not alter cell survival or behavior**
(A) Macrophages, labelled with an *mpeg1* promoter driving membrane YFP, display no difference in cell count or morphology when compared to cells with overexpressed cd44aTMICD under the *mpeg1* promotor. (B) Similarly, there is no noticeable change in appearance of xanthophore lineages overexpressing cd44aTMICD when compared to control fish. Scale bars represent 100µm (A, B).

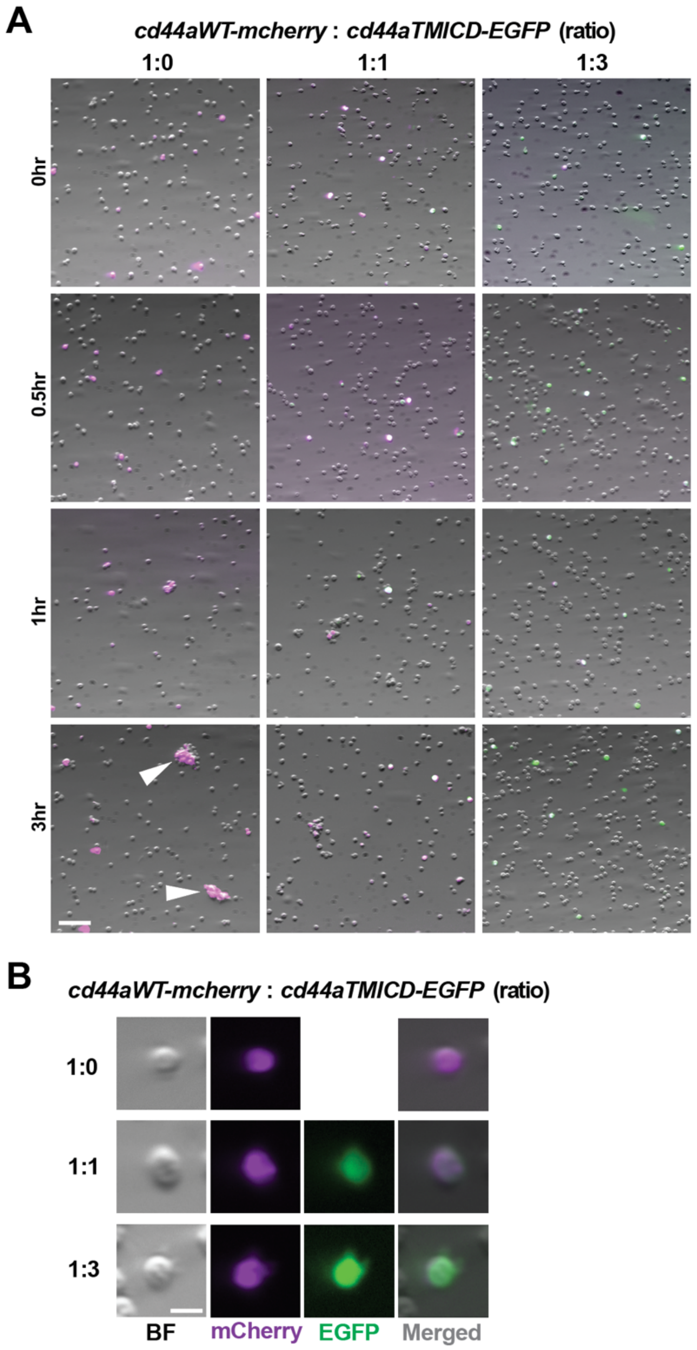


**Figure S4. Overexpressed CD44aTMICD interferes adhesive interaction mediated by wild type CD44a**

(A) S2 cells were transfected with constructs of cd44aWT-mCherry and cd44aTMICD-EGFP at various indicated ratios. Unlike the S2 cells transfected solely with cd44aWT-mCherry, those co-transfected with cd44aTMICD-EGFP exhibited a reduced capacity to form aggregates. Large cell aggregates were observed in cells expressing the wild type cd44a (white arrowheads). Co-transfected cells appeared white due to the co-expression of EGFP and mCherry (pseudo-colored in magenta), which are fused to either cd44aTMICD or cd44aWT. Cells expressing higher levels of EGFP in the 1:3 group displayed a greenish hue in white. (B) Single-cell resolution of transfected S2 cells at various indicated ratios. Note that CD44aWT-mCherry fusion protein expression was unaffected by co-expression with CD44aTMICD-EGFP. Quantifications can be found in Fig. 4C. Scale bar: 100μm (A), 20μm (B).

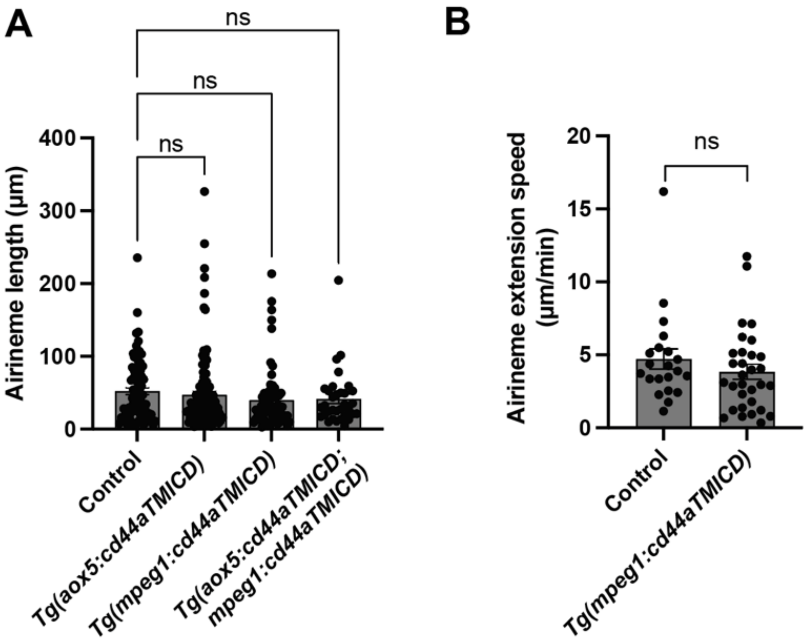


**Figure S5. Airineme length and speed remained consistent despite CD44a manipulations**
(A) There was no significant difference in airineme length in embryos where cd44aTMICD was overexpressed either in xanthophore-lineages, macrophages, or in both simultaneously, ($F_{(3, 288)}$=1.020, $P$=0.3843). (B) There was no statistically significant difference in airineme extension speed in embryos overexpressing cd44aTMICD in macrophages, $P$=0.2968, 3 embryos each. Statistical significance was assessed using a One-way ANOVA, followed by a Tukey's HSD post hoc test or a Student's t test. Error bars indicate mean ± SEM.

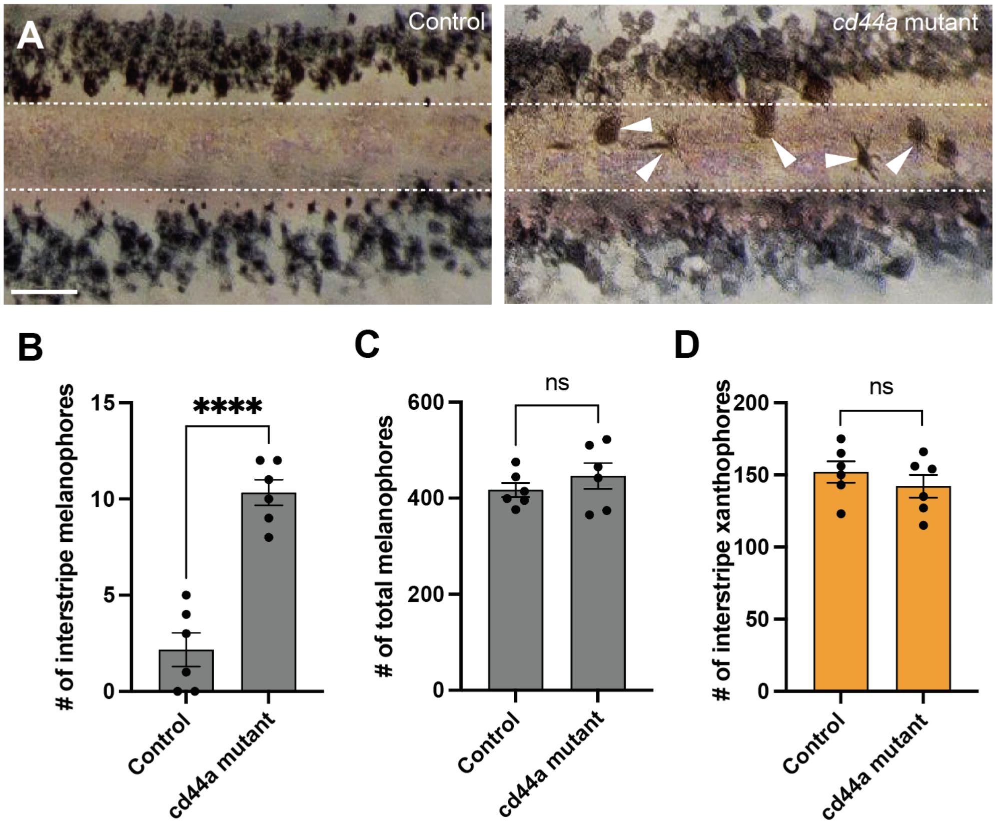


**Figure S6. Mutants of cd44 induced by CRISPR/Cas9 exhibit defective zebrafish pigment patterns**

(A) Melanophores failed to coalesce into stripe and remained in the interstripe zone (white arrowheads) in cd44 mutants. White dotted lines demarcate stripes and interstripe. (B) In cd44 mutant embryos, the count of interstripe melanophores was significantly higher, ($P$<0.0001, 12 embryos in total). (C) However, the total number of melanophores did not differ significantly between the experimental group and controls, ($P$=0.3732, 12 embryos in total). (D) The number of xanthophores in the interstripe did not differ significantly between the experimental group and controls (P=0.388, 12 embryos in total). Statistical significance was assessed using a Student's t test. Scale bars represent 200μm. Error bars indicate mean ± SEM.
